# Intrinsically disordered regions and RNA binding domains contribute to protein enrichment in biomolecular condensates in *Xenopus* oocytes

**DOI:** 10.1101/2023.11.10.566489

**Authors:** Liam C. O’Connell, Victoria Johnson, Anika K. Hutton, Jessica P. Otis, Anastasia C. Murthy, Mark C. Liang, Szu-Huan Wang, Nicolas L. Fawzi, Kimberly L. Mowry

## Abstract

Proteins containing both intrinsically disordered regions (IDRs) and RNA binding domains (RBDs) can phase separate *in vitro*, forming bodies similar to cellular biomolecular condensates. However, how IDR and RBD domains contribute to *in vivo* recruitment of proteins to biomolecular condensates remains poorly understood. Here, we analyzed the roles of IDRs and RBDs in L-bodies, biomolecular condensates present in *Xenopus* oocytes. We show that a cytoplasmic isoform of hnRNPAB, which contains two RBDs and an IDR, is highly enriched in L-bodies. While both of these domains contribute to hnRNPAB self-association and phase separation *in vitro* and mediate enrichment into L-bodies in oocytes, neither the RBDs nor the IDR replicate the localization of full-length hnRNPAB. Our results suggest a model where the additive effects of the IDR and RBDs regulate hnRNPAB partitioning into L-bodies. This model likely has widespread applications as proteins containing RBD and IDR domains are common biomolecular condensate residents.

## Introduction

Biomolecular condensates have emerged as a key feature of cellular architecture, serving to organize biomolecules such as RNAs and proteins into discreet bodies without a lipid membrane^1–3^. Biomolecular condensates are found in both the cytoplasm and the nucleus, and display a range of functions and biophysical properties^4^. The biophysical behavior of a biomolecular condensate tends to align with its function: transient structures like stress granules are more liquid-like and dynamic^5–7^, while biomolecular condensates that persist on the order of days to months, such as L-bodies, P-granules and germline granules, are less dynamic and more hydrogel- or solid-like in their physical state^8–11^.

*In vitro* studies performed with a variety of proteins and RNAs found in biomolecular condensates have elucidated many of their governing principles^12–14^. Prominent among these is the prevalence of intrinsically disordered regions (IDRs) in biomolecular condensate constituent proteins^13,15,16^. IDRs are regions of a protein that display low degrees of sequence complexity, and consequently are unable to form stable secondary or tertiary structures, remaining more or less unfolded^5^. Many studies have shown that proteins containing IDRs phase separate *in vitro* via protein-protein interactions, forming bodies that display properties similar to biomolecular condensates^7,12,17–19^. Many IDR-containing proteins also have RNA binding domains (RBDs). Studies have consistently shown that addition of RNA to RNA binding proteins (RBPs) *in vitro* facilitates the formation of phase separated bodies, indicating a role for RNA-protein interactions in their formation in addition to IDR interactions^19–21^. This is notable given that many biomolecular condensates found in cells are composed of RNA and protein, suggesting that both protein-protein and RNA-protein interactions occur in *in vivo* phase-separated bodies^22^. However, while the mechanics of *in vitro* phase separated bodies have been well-studied, how these principles translate to more complex heterogeneous biomolecular condensates in cells remains less clear.

The *Xenopus laevis* oocyte organizes its cytoplasm in part through RNA localization, which specifies germ layer patterning during embryogenesis^23,24^. mRNAs encoding morphogens such as the TGF-β growth factor *vg1* are transported to the vegetal cortex during oogenesis via large cytoplasmic granules containing both RNA and protein^25^. Work from our laboratory discovered that these RNP granules are novel cytoplasmic biomolecular condensates termed Localization bodies, or L-bodies ^26^. Localizing RNAs are highly enriched in L-bodies and form a non-dynamic scaffold within them, in contrast to L-body constituent proteins, which display a range of dynamics, from moderate to high^26^. As with several other biomolecular condensates, the L-body proteome is enriched for proteins containing RBDs and IDRs, with many proteins containing both^26^. While RBDs and IDRs have been shown to be integral to phase separation *in vitro*, their relative importance to protein enrichment and dynamics in biomolecular condensates *in vivo* remains less clear, particularly in the context of proteins that contain both types of domains.

L-bodies present a unique opportunity to study how proteins associate with long-lived biomolecular condensates, as other experimentally tractable biomolecular condensates such as stress granules tend to be short-lived^27^. As such, previous studies have often focused on protein recruitment via de novo assembly of biomolecular condensate^28–30^, whereas L-bodies allow us to investigate how proteins are recruited to extant biomolecular condensates.

In this work we investigated how RBD and IDR domains regulate protein enrichment and dynamics within L-bodies. We focused on the *Xenopus* RBP heterogenous nuclear ribonucleoprotein AB (hnRNPAB) due to its known localization to L-bodies, association with L-body constituent mRNA *vg1*, and structure, which consists of two RBDs and C-terminal IDR ^31,32^.

We found that hnRNPAB X2, a novel cytoplasmic splice isoform of hnRNPAB, lacks the C-terminal nuclear localization signal found in the canonical form of hnRNPAB and is highly enriched in L-bodies. While both the hnRNPAB X2 RBDs and IDR are sufficient individually for L-bodies enrichment, enrichment was lower than for the full-length protein, indicating that the two domains together are necessary to replicate enrichment of hnRNPAB X2 in phase separated L-bodies. Consistent with this, both the RBD and the IDR domains phase separate *in vitro* and the dynamic behavior of hnRNPAB *in vivo* is dependent on both domains. While hnRNPAB X2 is fully dynamic within L-bodies, the RBD and IDR of hnRNPAB X2 are each more dynamic than the full-length protein. Moreover, the addition of the hnRNPAB X2 IDR to PTBP3, an L-body RBP that contains four RBDs and lacks an IDR, reduces the dynamics of the chimeric protein in L-bodies. Taken together, our results suggest an additive model for understanding how functional domains act to regulate protein enrichment and dynamics within biomolecular condensates such as L-bodies.

## Results

### hnRNPAB X2 is a novel cytoplasmic isoform of hnRNPAB

To investigate how RBDs and IDRs regulate protein enrichment and dynamics in biomolecular condensates we focused on hnRNPAB, a constituent of L-bodies that contains two RBDs and an IDR (Figure S1a). *vg1* mRNA is transported to the vegetal cortex of *Xenopus* oocytes during stages II-III of oogenesis in L-bodies in a process that can be visualized by imaging a fluorescently-labeled localization element (LE) derived from sequences from the 3′ UTR of *vg1* mRNA^33,34^. Analysis of the distribution of endogenous hnRNPAB by immunostaining in stage II-III oocytes showed that hnRNPAB is present in both the nucleus and cytoplasm, with notable enrichment to the vegetal cortex and L-bodies as demonstrated by colocalization with *LE* RNA (Figure 1a). However, expression of mCherry-tagged hnRNPAB in stage II-III oocytes following microinjection of a cloned, tagged mRNA showed predominantly nuclear localization (Figure 1b). This led us to ask whether an endogenous hnRNPAB splice isoform represented the cytoplasmic protein seen for the endogenous protein by immunofluorescence (IF). We identified a computationally predicted splice isoform, hnRNPAB X2 (Figure 1c), and successfully cloned a cDNA from stage II oocyte RNA corresponding to the predicted sequence. IF of mCherry-tagged hnRNPAB X2 (Figure 1e) showed exclusively cytoplasmic signal with enrichment in L-bodies and the vegetal cortex similar to that observed in the cytoplasm by IF for endogenous hnRNPAB (Figure 1a). These results indicate that the distribution of endogenous hnRNPAB observed by IF (Figure 1a) represents the combined distribution of two isoforms, with the hnRNPAB X2 splice isoform localized in the cytoplasm and canonical hnRNPAB in the nucleus.

**Figure 1:**
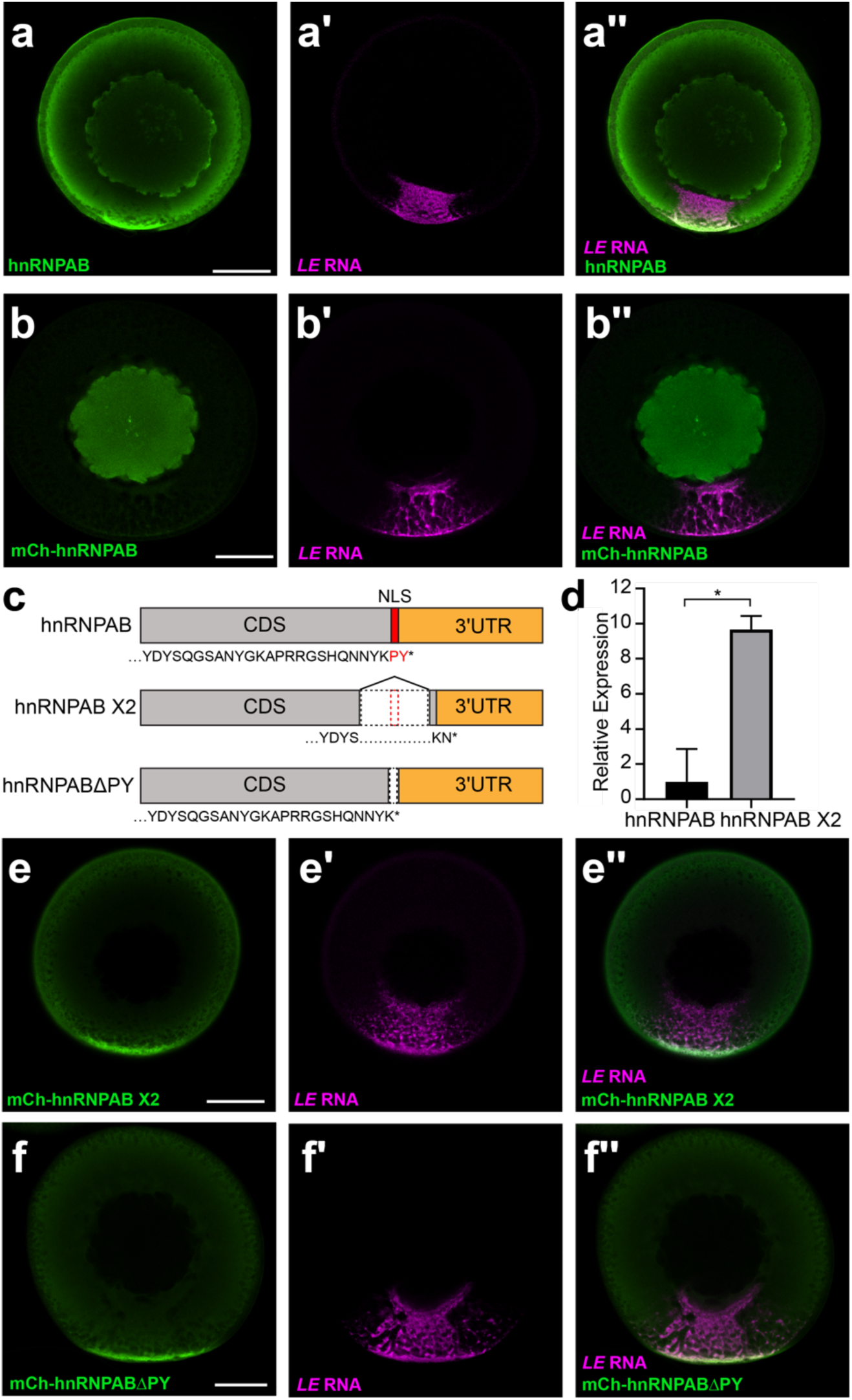
hnRNPAB X2 is a novel cytoplasmic splice isoform of hnRNPAB that is enriched in L-bodies. **(a)** Stage II oocytes were microinjected with Cy5-labeled *LE* RNA (magenta, a′) and immunostained for endogenous hnRNPAB (green, a) using antibodies raised against *Xenopus* hnRNPAB^32^. The overlap is shown in a″. **(b)** Cy5-labeled *LE* RNA (magenta, b′) was microinjected into stage II oocytes expressing mCherry-tagged canonical hnRNPAB, as detected by immunostaining with anti-mCherry (green, b). The overlap is shown in b″. **(c)** Schematics of canonical hnRNPAB, hnRNPAB X2 and hnRNPABΔPY. C-terminal sequence is shown below each, with the PY NLS indicated in red, the protein coding sequence (CDS) in gray, and the 3′UTR shown in orange. **(d)** RNA isolated from stage II oocytes was used to measure the relative expression of hnRNPAB X2 compared to canonical hnRNPAB by qPCR. ΔCt values were calculated normalizing to refence gene *vg1* (* indicates p<0.05 by T-test). Error bars represent standard deviation from the mean, *n*=3. **(e)** Cy5-labeled *LE* RNA (magenta, e′) was microinjected into stage II oocytes expressing mCherry-tagged hnRNPAB X2, as detected by immunostaining with anti-mCherry (green, e). The overlap is shown in e″. **(f)** Cy5-labeled *LE* RNA (magenta, e′) was microinjected into stage II oocytes expressing mCherry-tagged hnRNPABΔPY, as detected by immunostaining with anti-mCherry (green, e). The overlap is shown in e″. Confocal sections (a-b, e-f) are shown with the vegetal hemisphere at the bottom; scale bars=100µm.

Pre-mRNA processing of the hnRNPAB X2 isoform splices out nucleotides encoding the C-terminal region of the protein (Figure 1c) which contains a putative PY nuclear localization sequence (NLS)^35^; related proteins such as JKTBP and hnRNPD have a NLS in this region, which we hypothesized might be ablated in hnRNPAB X2. To test this, we ablated the putative PY NLS in canonical hnRNPAB (construct hnRNPABΔPY; Figure 1c) by deleting the two most C-terminal amino acids (PY). Expression and imaging of this construct (Figure 1f) showed that deletion of the C-terminal PY residues is sufficient to recapitulate hnRNPAB X2-like cytoplasmic localization (Figure 1e-f), suggesting that hnRNPAB X2 represents an isoform of hnRNPAB that is targeted to the cytoplasm through the loss of a C-terminal NLS. Indeed, qPCR carried out with primers specific to canonical hnRNPAB or hnRNPAB X2 show that hnRNPAB X2 is ∼10-fold more highly expressed than the canonical splice isoform in stage II-III oocytes (Figure 1d).

### Both the RBDs and the IDR contribute to hnRNPAB X2 L-body localization

We next investigated the roles of the RBD and IDR domains of hnRNPAB X2 in its enrichment in L-bodies. We first analyzed hnRNPAB X2 via Prion Like Amino Acid Composition (PLAAC)^36^ and found that the IDR has significant predicted Prion-like character (Figure S1a). Based on the PLAAC analysis and structured domain annotation, we generated two mCherry (mCh) tagged constructs termed RBD and IDR, which each isolated their respective functional domains (Figure 2a). The RBD construct contains two RNA recognition motif (RRM) class RNA-binding domains, and the IDR construct contains the prion-like IDR (Figure 2a). In the case of stress granule localization of the human RBP Fused in Sarcoma (FUS), that contains a prion-like domain (very much like that of hnRNPAB) and RNA-binding domains (RRM, zinc finger, as well as disordered RGG motifs in the case of FUS), the RBDs but not the prion-like domains are sufficient for, and contribute to, incorporation in stress granules^37^. Therefore, we hypothesized that the hnRNPAB X2 RBD is required for recruitment to L-bodies and the IDR is dispensable. Expression of the mCh tagged RBD and IDR constructs in oocytes co-microinjected with fluorescent *LE* RNA indicated that both were enriched in L-bodies relative to free mCh (Figure 2b-e), and were expressed at levels similar to full-length hnRNPAB X2 (Figure S2). To evaluate L-body enrichment, images of mCh-hnRNPAB X2, mCh-RBD, mCh-IDR, and mCh alone were blinded and scored on a scale of 0-3, where 0 indicated no L-body colocalization and 3 indicated strong colocalization with L-body localizing RNA (Figure 2f). While mCh-RBD and mCh-IDR were significantly enriched in L-bodies relative to free mCh, hnRNPAB X2 was significantly more enriched than either of the isolated domains, indicating that neither domain alone is sufficient to mimic hnRNPAB X2 localization. Therefore, it is likely that hnRNPAB X2 localizes to L-bodies due to a combination of interactions by both the RBD and IDR domains.

**Figure 2:**
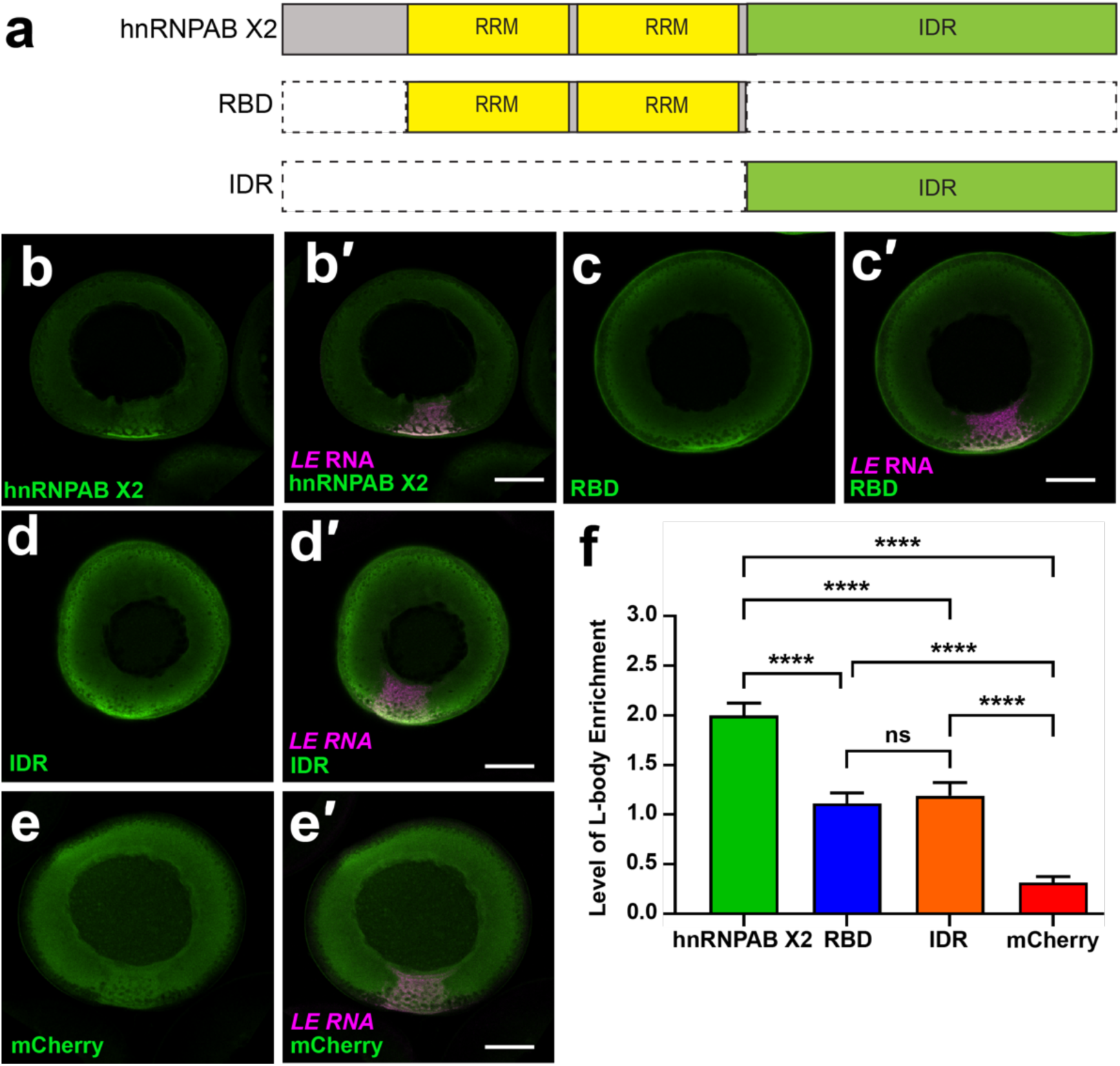
hnRNPAB X2 localizes to L-bodies through its RBD and IDR. **(a)** Schematics of domain constructs. **(b-e)** Stage II oocytes expressing (b) mCh-hnRNPAB X2, (c) mCh-RBD, (d) mCh-IDR, or (e) free mCh (green; detected by anti-mCherry IF) were co-microinjected with Cy5 *LE* RNA (magenta, b′-e′) to label L-bodies. Colocalization (white) is shown in the merged confocal images (b′-e′). Scale bars=100μm. **(f)** Scoring of L-body enrichment (*n*=21 oocytes) where 0 indicates no enrichment of the protein in L-bodies (as marked by *LE* RNA), 1 indicates modest enrichment, 2 indicates moderate enrichment and 3 indicates strong enrichment. Shown are levels of L-body enrichment for hnRNPAB X2 (green), RBD (blue), IDR (orange), and mCherry (red). *n*=21 oocytes, error bars represent standard error of the mean, **** indicates *p* <0.001and ns indicates *p* >0.05. Statistics shown are an Ordinary one-way ANOVA followed by Tukey’s multiple comparisons.

### The hnRNPAB X2 IDR and RBD domains both phase separate *in vitro*

Biomolecular condensates have been characterized to form through phase separation^38^. We used an *in vitro* approach to test if hnRNPAB X2 or its domains phase separate: differential interference microscopy on samples of recombinant hnRNPAB X2 purified as fusions with maltose binding protein (MBP) solubility tags. In the presence of crowding agent (10% PEG), full-length hnRNPAB X2 formed irregularly shaped assemblies consistent with static structures (Figure 3a). Conversely, the RBD and IDR domains both formed round droplets that flow, fuse, and round up, consistent with dynamic liquid assemblies (Figure 3b-c). Notably, a control protein (MBP alone) did not show phase separation (Figure 3d). These *in vitro* phase-separated assemblies are similar in the presence of RNA (Figure S3). Taken together, these results indicate that, independent of the presence of RNA, full-length hnRNPAB X2 is prone to self-assemble and both the RBD and IDR domains are sufficient to generate liquid phase separated droplets *in vitro*.

**Figure 3:**
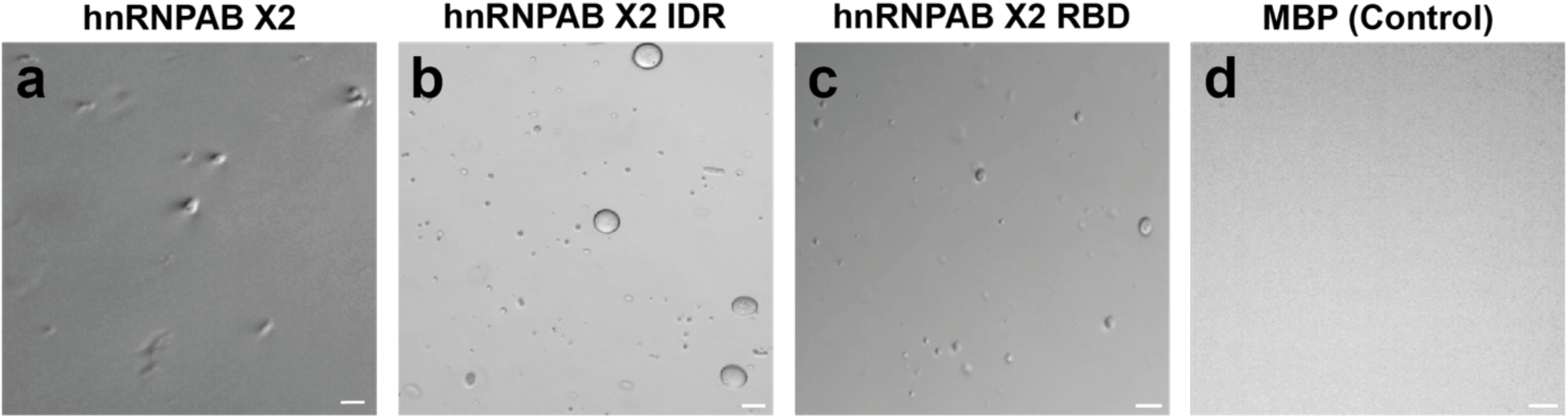
hnRNPAB X2 and its domains self-assemble and phase separate *in vitro*. DIC micrographs of 50 µM MBP-fusions of (**a**) full-length hnRNPA2 X2, **(b)** IDR, (**c**) RBD, or (**d**) control (free MBP) proteins in 20 mM NaPi (pH 7.4), 150 mM NaCl, and 10% PEG. Irregularly shaped assemblies are observed for hnRNPA2 X2 full-length (a), while round droplets consistent with liquid-liquid phase separation are observed for hnRNPAB X2 IDR and RBD (b-c) domains. No phase separation is observed for MBP alone (control, d). Images are representative from two biological replicates (with independently expressed and purified protein). Scale bars=20 μm.

### hnRNPAB X2 is highly dynamic in L-bodies

We next sought to determine the dynamics of hnRNPAB X2 in L-bodies and how the IDR and RBD domains regulate hnRNPAB X2 dynamics in L-bodies *in vivo*. Proteins containing IDRs, including hnRNPAB, have been shown to dynamically associate with biomolecular condensates like L-bodies^39^. Therefore, we hypothesized that the RBDs may form stable RNA interactions and therefore be less dynamic than the full-length protein and that the IDR modulates the dynamic physical association of hnRNPAB X2 within L-bodies. We assessed the dynamics of mCh-tagged hnRNPAB X2 by fluorescence recovery after photobleaching (FRAP), defining the regions of interest (ROIs) such that L-bodies were only partially bleached, to allow for recovery both from within the L-body and from the surrounding cytoplasm (Figure 4a). Despite its enrichment to L-bodies, hnRNPAB X2 is completely dynamic within them, with a mobile fraction of 100% (Figure 4b). We then tested whether the dynamics of hnRNPAB X2 were governed by the IDR domain, reasoning that the RBDs alone might display less dynamic behavior, due to potential interactions with the non-dynamic RNA phase. We tested the mCh-RBD and mCh-IDR constructs by FRAP and found that both the RBD and the IDR display mobile fractions of 100% identical to the full-length protein (Figure 4c), contradicting the idea that the RBD would display lower dynamics. Notably, both mCh-RBD and mCh-IDR display significantly shorter t_1/2_ measurements (8 and 9 seconds, respectively) than the full-length protein (31 seconds) (Figure 4d). These FRAP experiments suggests that, similar to protein enrichment in L-bodies, the dynamic behavior of hnRNPAB X2 is influenced by both the IDR and RBD domains, and that neither type of domain alone is sufficient to rescue the behavior of the full-length protein.

**Figure 4:**
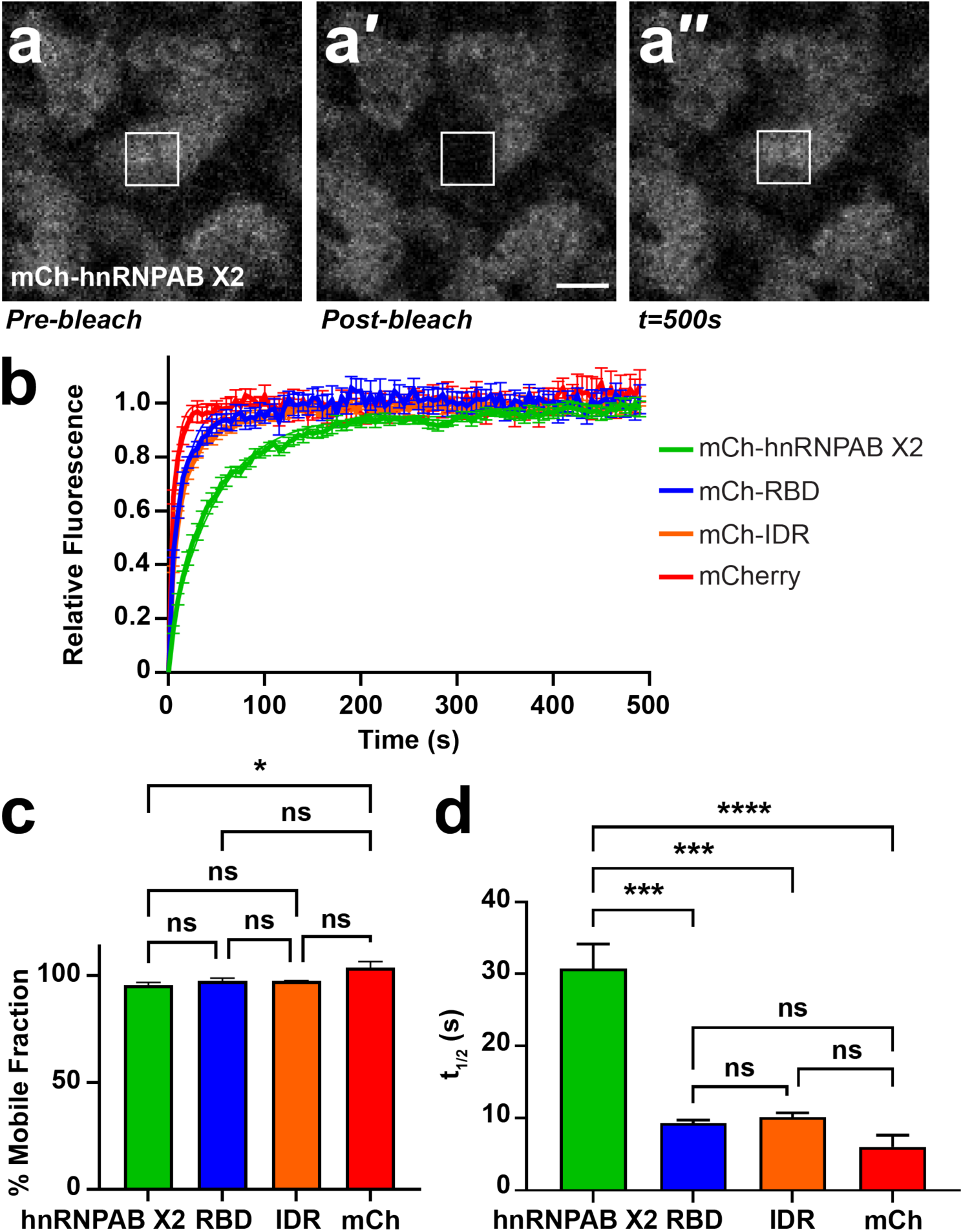
hnRNPAB X2 dynamically associates with L-bodies via its RBD and IDR. **(a)** Shown is an image of the vegetal cytoplasm of a stage II oocyte microinjected with mCh-hnRNPAB X2. FRAP was conducted such that an individual L-body was partially bleached (a′), to allow for recovery (a″) both from within the L-body and from its environment. The 10µm^2^ ROI is indicated by a white box; scale bar=10μm. **(b)** Stage II oocytes were microinjected with mCherry (mCh), mCh-hnRNPAB X2, mCh-RBD or mCh-IDR RNA to express the mCh-tagged proteins, along with Cy5 *LE* RNA to mark L-bodies. Normalized FRAP recovery curves are shown (*n*=21 oocytes per fusion protein); error bars represent SEM. Measurements were taken at 5 second intervals over 100 iterations. **(c)** Plateau values (% mobile fraction) for each construct are shown (ns indicates not significant, * indicates p<0.05). Error bars represent SEM. Statistics shown are an Ordinary one-way ANOVA with Tukey’s multiple comparisons. **(d)** T_1/2_ measurements are shown for each construct, measured as the average time at which the construct recovers half of its plateau value (ns indicates not significant, *** indicates p<0.001, **** indicates p<0.0001). Error bars represent SEM. Statistics shown are an Ordinary one-way ANOVA with Tukey’s multiple comparisons.

### The hnRNPAB X2 IDR stabilizes proteins in L-bodies

Given that the IDR of hnRNPAB X2 localizes to, and is fully dynamic within, L-bodies, we next asked if the dynamic behavior of hnRNPAB X2 could dominate a more stable L-body protein. Work from our laboratory has recently shown that the RBP PTBP3 is a core component of L-bodies with moderate dynamics (mobile fraction ∼48%)^40^. PTBP3 contains four well-folded RRM domains but no discernable IDR (Figure S1B). We generated a mCh-tagged construct of PTBP3 fused with the hnRNPAB X2 IDR, termed PTBP3+IDR (Figure 5a). Expression and imaging of mCh-PTBP3+IDR showed localization to L-bodies similar to wild type PTBP3 (Figure 5b)^40^. However, upon assaying PTBP3+IDR by FRAP, we found that the addition of the hnRNPAB X2 IDR decreased the dynamics of PTBP3 in L-bodies, as the chimeric protein displayed a mobile fraction of only 15% (Figure 5c-d), indicating that the hnRNPAB X2 IDR domain can stabilize proteins within L-bodies.

**Figure 5:**
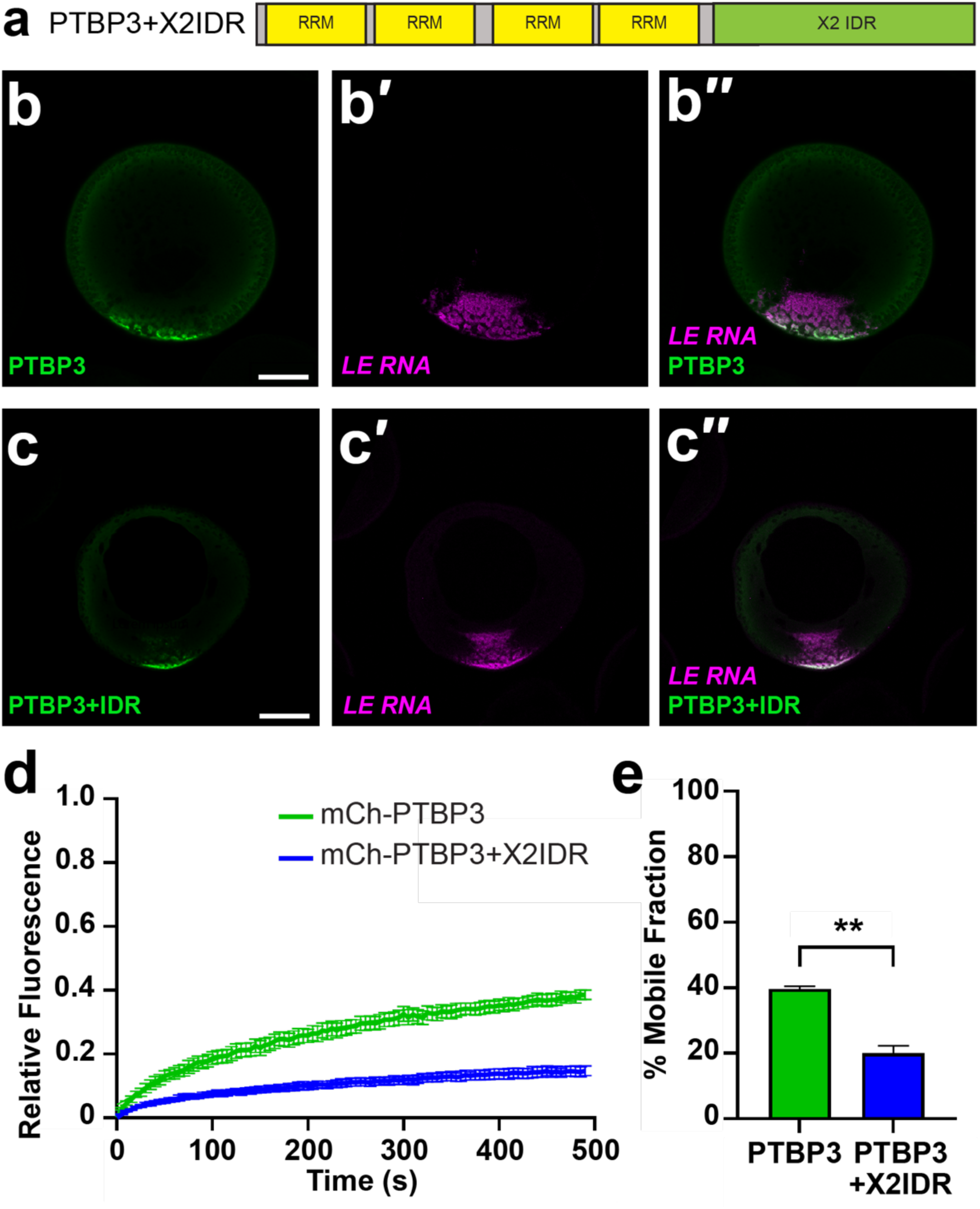
The hnRNPAB X2 IDR acts to stabilize proteins in L-bodies. **(a)** The hnRNPAB X2 IDR was fused to the C-terminus of PTBP3 (PTBP3+X2IDR) and tagged with mCherry (mCh). **(b)** Stage II oocytes were microinjected with mCh-PTBP3 RNA to express the encoded protein (green, detected by anti-mCh IF) along with Cy5 *LE* RNA to label L-bodies (magenta, b′). The merge is shown in b″; scale bar=100μm. **(c)** Stage II oocytes were microinjected with mCh-PTBP3+IDR to express the encoded protein (green, detected by anti-mCh IF) along with Cy5 *LE* RNA (magenta, c′) to label L-bodies. The merge is shown in c″; scale bar=100μm. **(d)** Stage II oocytes were microinjected with RNA encoding PTBP3 or PTBP3+X2IDR, along with Cy5 *LE* RNA to label L-bodies. Normalized FRAP recovery curves are shown (*n*=21 oocytes); error bars represent SEM. Measurements were taken at 5 second intervals over 100 iterations. **(e)** Average percent mobile fractions for the two constructs are shown. Error bars represent SEM; ** indicates *p*<0.01, ns indicates not significant (*p*>0.05). Statistics shown are an Ordinary one-way ANOVA with Tukey’s multiple comparisons.

**Figure 6:**
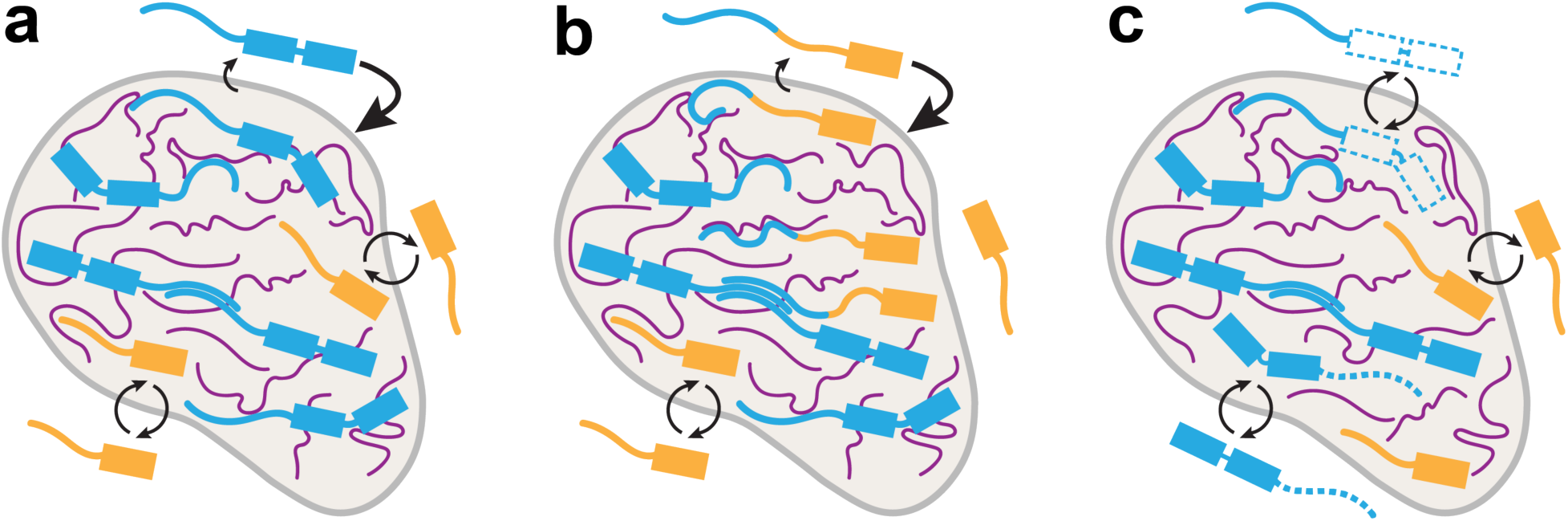
A model for how RBDs and IDRs additively determine a protein’s localization (hnRNPAB) to and dynamics within L-bodies. **(a)** While RNA (magenta) is highly stable within L-bodies (grey), L-body associated proteins (blue, gold) dynamically associate with the RNA and other proteins. The strength of these associations determines the degree of their restriction to L-bodies: proteins that interact strongly with L-bodies (blue) show a greater degree of enrichment to L-bodies and lower degrees of dynamic activity, while proteins that interact more weakly (gold) are less enriched and more dynamic. **(b-c)** The association of a protein with L-bodies can be tuned by modifying its interaction domains. Additional interaction domains (b) may stabilize a protein within L-bodies, while removal of interaction domains (c) may serve to destabilize a protein within L-bodies. This principle holds true for both RBDs (represented by boxes) and IDRs (represented by lines).

## Discussion

We investigated how a protein containing RBDs and an IDR localizes to a biomolecular condensate. The RBD- and IDR-containing protein hnRNPAB X2 is both highly enriched in L-body biomolecular condensates and extremely dynamic within them. Interestingly, the nuclear form of hnRNPAB, which contains the same architecture with the addition of only a short nuclear localization sequence, is not assembled into granules. This difference may be due to the high concentration of RNA throughout the nucleus which can prevent phase separation of RBPs^38^ or due to the lack of L-body-specific components that would be needed to recruit hnRNPAB to other nuclear bodies. We also probed the contribution of each domain to hnRNPAB X2 behavior within L-bodies. The isolated hnRNPAB X2 RBD and IDR domains are sufficient for localization to L-bodies, but to a lesser degree than the full-length protein. *In vitro*, full-length hnRNPAB X2 protein forms assemblies that appear rigid and static while both the IDR and RBD domains display more liquid-like phase separation. Hence, both domains contribute to hnRNPAB X2 self-interactions and phase separation *in vitro*. Furthermore, this self-interaction does not require RNA, similar to other RBPs such as FUS, which can phase separate readily without RNA and whose disordered domain phase separates but does not interact with RNA^41^. Both the RBD and IDR domains of hnRNPAB X2 also affect protein dynamics in L-bodies: as assessed by FRAP both the RBD and IDR displayed shorter recovery half-times than the full-length protein, indicating that interactions anchoring the domains to L-bodies are reduced with the removal of the other domain.

Taken together, our results suggest an additive model in which *in vivo* enrichment of RBPs like hnRNPAB X2 that contain phase separation prone IDRs is determined by the sum of its interactions with L-body constituents, independent of the types of forces governing those interactions. Similarly, this model also suggests that RBP dynamics within L-bodies is determined by the combined effects of the interaction domains found in a protein. We had hypothesized that increasing an RBP’s association with the dynamic protein layer of L-bodies (such as by adding the hnRNPAB X2 IDR to PTBP3, which is known to bind directly to *LE* RNA) would cause it to be more dynamic, but found instead that adding the IDR causes the protein to become less dynamic and more stably associated with L-bodies. This indicates that this additive model is likely agnostic of the manner in which a protein interacts with L-bodies; whether it binds the stable RNA directly as PTBP3 does or interacts with the dynamic protein layer as the hnRNPAB X2 IDR might, these interactions serve to enrich and stabilize the protein in L-bodies.

Indeed, it is possible that this model extends beyond proteins and that the stable RNA phase we observe in L-bodies^10,40^ is in fact an extreme end of this sliding scale of interaction strength. We previously showed^40^ that mutating *LE* RNA such that it could no longer bind two sequence-specific RBPs abolished its ability to localize to L-bodies and caused a sharp increase in its percent mobile fraction (from ∼5% to ∼60%). This linkage in *LE* RNA between a reduction in interaction strength, reduced localization, and increased dynamics is similar to what we have found with hnRNPAB X2 and illustrates what may be fundamental principles underlying the behavior of L-body constituents. In addition, our results, which indicate that IDRs play a role in protein recruitment to stable, long-lived biomolecular condensates, mirror those found in other systems^42^, suggesting that while a biomolecular condensate may mature over time, the fundamental physical properties governing protein recruitment to condensates are likely to remain constant.

## Methods

### Oocyte Isolation and Culture

All animal experiments were approved by the Brown University Institutional Animal Care and Use Committee. Oocytes were harvested from wild type *Xenopus laevis* females (Nasco, catalog # LM00535MX). Oocytes were enzymatically defolliculated in 3 mg/ml collagenase (Sigma) followed by washes in MBSH (88mM NaCl, 1mM KCl, 2.4mM NaHCO_3_, 0.82mM MgSO_4_, 0.33mM CaCl_2_, 0.33mM Ca(NO_3_)_2_, 10mM HEPES pH7.6). Stage II-III oocytes were cultured at 18°C in XOCM [50% Leibovitz L-15 (Thermofisher), 15mM HEPES (pH 7.6), 1mg/mL insulin, 50U/mL nystatin, 100U/mL penicillin/streptomycin, 0.1mg/mL gentamicin].

### Cloning

Total RNA from stage II-III oocytes was extracted by TRIZOL. cDNA was generated via the iScript cDNA synthesis kit. Primers specific to hnRNPAB X2 (Table S3) were used to amplify hnRNPAB X2 from cDNA using Phusion PCR master mix which was then cloned into pSP64TNRLMCS:mCh^10^ using Gibson Assembly master mix (New England Biolabs) to generate pSP64:mCh-hnRNPAB-X2. Table S3 shows the primer sequences used to generate pSP64:mCh-RBD-X2, pSP64:mCh-IDR-X2, and pSP64:mCh-hnRNPABΔPY using pSP64:mCh-hnRNPAB-X2 as a template for PCR, followed by cloning into pSP64TNRLMCS:mCh as described above. Plasmids for expression of hnRNPAB and its isolated domains with a N-terminal maltose-binding protein tag were generated by PCR using primers (Table S3) specific to the appropriate sequence region with NdeI and XhoI sites for subcloning into the pTHMT vector^41^.

### RNA transcription and microinjection

Fluorescently labeled *vg1 LE* RNA (Gautreau et. al., 1997) was generated from linearized pSP73-2x135 using the MEGAscript T7 transcription kit (Ambion) in the presence of 250 nM Cy^TM^5-UTP. mCherry (mCh)-tagged protein-coding mRNAs were generated using the mMessage Machine transcription kit from the following linearized plasmids: pSP64:mCh-hnRNPAB-X2, pSP64:mCh-RBD-X2, pSP64:mCh-IDR-X2, pSP64:mCh, pSP64:mCh-hnRNPAB-FL, pSP64:mCh-hnRNPABΔPY, pSP64:mCh-PTBP3 and pSP64:mCh-PTBP3+IDR. mRNAs and Cy-labeled RNAs were co-microinjected at 500 nM at a volume of 2 nL per oocyte. Following injection, oocytes were cultured for ∼48 hours at 18°C in XOCM. Unlabeled *Xenopus* β-globin RNA was transcribed with the MEGAscript T7 kit (ThermoFisher) using a PCR product generated from pSP64:XBM^43^ (see Table S3 for primers).

### Whole-mount immunofluorescence

Oocytes were washed in PBS to remove XOCM and then fixed for 1 hour at room temperature in PEMT-FA (80 mM PIPES pH6.8, 1 mM MgCl_2_, 5 mM EGTA, 0.2% Triton X-100, 3.7% formaldehyde). Fixed oocytes were washed in PBT (137 mM NaCl, 2.7 mM KCl, 10 mM Na2HPO4, 1.8 mM KH2PO4, 0.2% BSA, 0.1% Triton X-100) 3 times for 15 minutes each, then blocked in PBT+ (PBT supplemented with 2% normal goat serum, 2% BSA) for 2 hours at room temperature. Oocytes were then incubated overnight at 4° C in a 1:500 dilution of primary antibody (listed in Table S2) in PBT+. Following incubation, oocytes were washed three times for two hours each in PBT, then incubated overnight at 4° C in a 1:1000 dilution of secondary antibody (Thermofisher goat anti-rabbit AF546, 11010) in PBT+. Oocytes were then washed three times for two hours each in PBT. Following the washes, oocytes were dehydrated in anhydrous methanol and stored at −20° C until imaging. Immediately prior to imaging, oocytes were cleared in BABB solution (1:2 benzyl alcohol:benzyl benzoate). Oocytes were imaged on an inverted Olympus FV3000 confocal microscope using 20x UPlan Super Apochromat objective (air, NA=0.75) and 30x UPlan Super Apochromat objective (silicon oil, NA=1.05) using GaAsP detectors. To score localization, images were blinded and scored on a scale of 0-3, where 0 indicated no L-body colocalization and 3 indicated strong colocalization with L-body localizing RNA.

### *In vitro* expression and purification of hnRNPAB X2 full length protein, IDR, and RBD domains

N-terminally MBP tagged (pTHMT) hnRNPAB X2 full-length isoform, IDR and RBD domains, and MBP (control) were expressed in *E. coli* BL21 star (DE3) cells (C600003; Thermofisher Scientific). Bacterial cultures were grown to an optical density at 600 nm of 0.7-0.9 before induction with 1 mM isopropyl-β-D-1-thiogalactopyranoside (IPTG) for 4 hours at 37° C. Cell pellets were harvested by centrifugation and stored at −80°C. Cells were resuspended in ∼40 mL of 20 mM Tris, 500 mM NaCl, 10 mM Imidazole (pH 8.0), with one Roche EDTA-free protease inhibitor tablet (11697498001; Sigma Aldrich) per ∼4 g cell pellet, and lysed using the Avestin Emulsiflex C3 (Ottawa, Ontario, Canada). The lysate was cleared by centrifugation at 47,850 x g (20,000 rpm) for 50 min at 4°C, filtered using a 0.2 µm syringe filter, and loaded onto a HisTrap HP 5 ml column (17524701; Cytiva, Marlborough, MA, USA). The protein was eluted with a gradient from 10 to 300 mM imidazole in 20 mM Tris, 500 mM NaCl, pH 8.0. Fractions containing MBP-tagged hnRNPAB X2 full length isoform were loaded onto a HiLoad 26/600 Superdex 200 pg column (28-9893-36; Cytiva) equilibrated in 20 mM Tris, 150 mM NaCl, and 1 mM dithiothreitol (DTT). Fractions containing MBP-tagged IDR, -RBD, or MBP alone were loaded onto a HiLoad 26/600 Superdex 75 pg (28-9893-34; Cytiva) equilibrated in 20 mM Tris and 150 mM NaCl. Fractions with high purity were identified by SDS-PAGE and concentrated using a centrifugation filter with a 10 kDa cutoff (ACS501024; MilliporeSigma, Burlington, MA, USA). MBP-tagged hnRNPAB X2 full length isoform, the IDR and RBD domains, and MBP were flash frozen in liquid nitrogen in 20 mM Tris and 150 mM NaCl and stored at −80 °C.

### Differential interference contrast microscopy

Droplets were prepared in the purification buffer at 50 µM protein concentration for MBP-tagged hnRNPAB X2 full length isoform, and the IDR and RBD domains in 20 mM NaPi, pH 7.4, 150 mM NaCl, in the absence and presence of 0.25 mg/mL *Xenopus* β-globin RNA with the addition of 10% PEG to induce phase separation, analogous to recent protocols for other RNA-binding proteins^44^. After inducing phase separation in microcentrifuge tubes, the samples were incubated at room temperature for ∼30 min before visualization. Samples were spotted onto a glass coverslip and droplet formation was evaluated by imaging with differential interference contrast (DIC) on a Ti2-E Fluorescence microscope (Nikon). Images are representative from two biological replicates (with independently expressed and purified protein).

### FRAP

Stage II oocytes were microinjected with 2 nL of 500 nM mRNA encoding mCherry-tagged constructs and 250 nM Cy5-UTP labeled *LE* RNA to mark L-bodies. Oocytes were cultured in XOCM at 18° C for ∼48 hours post-injection. A 10 μm^2^ ROI was bleached using a 488 nm laser at 100% for 2 seconds. Recovery of the fluorescence was measured at 5 second increments over 100 iterations. Seven oocytes per biological replicate (21 oocytes total per protein construct) were analyzed. Fluorescence measurements were corrected based on background fluorescence and nonspecific photobleaching, as previously described^40,45,46^. Data were analyzed via one phase non-linear regression analysis using GraphPad Prism 9. Statistics shown in the main text for the percent immobile fractions are an ordinary one-way ANOVA with Tukey’s multiple comparison of the one phase association data.

### qPCR

One hundred stage II oocytes per biological replicate were washed in PBS and homogenized in Trizol to extract total oocyte RNA. Total oocyte RNA was reverse transcribed using the iScript cDNA synthesis kit (BioRad). qPCR was performed using SybrGreen Powerup Master mix (ThermoFisher, A25742) per the manufacturer’s protocol for *vg1*, canonical *hnRNPAB* and *hnRNPAB X2* RNAs, using the primers shown in Table S3. ΔCt values were calculated^47^ normalizing to *vg1* as a refence gene, and compared by T-test.

## Data Availability

All data supporting the findings of this study are available within the paper and its Supplementary Information.

## Acknowledgements

We thank Juan Alfonzo for comments on the manuscript. We thank Kevin Czaplinski for gifting the 40-LoVe (*Xenopus* hnRNPAB) antibody. This work was funded by R01GM071049 from the NIH to KLM and by R01GM147677 from the NIH to NLF.

## Author Contributions

LCO and KLM conceptualized the study and contributed to the review and editing of the manuscript. KLM carried out supervision, acquired funding, contributed to visualization, and editing of the manuscript. LCO carried out methodology development, investigation, formal analysis, visualization, and writing of the original draft. VJ carried out investigation, formal analysis, and visualization. AKH carried out investigation, formal analysis, and visualization. JPO carried out investigation, formal analysis, and visualization. ACM carried out methodology development and investigation. MCL carried out methodology development and investigation. S-HW carried out investigation. NLF carried out supervision, acquired funding, contributed to visualization, and editing of the manuscript.

