## Supplemental Information for "Intrinsically disordered regions and RNA binding domains contribute to protein enrichment in biomolecular condensates in *Xenopus* oocytes"

### Supplementary Information

O’Connell *et al.*

| | |
|---|---|
| Figure S1 | page 2 |
| Figure S2 | page 3 |
| Figure S3 | page 4 |
| Table S1 | page 5 |
| Table S2 | page 5 |
| Table S3 | pages 5-6 |
| References | page 7 |

### Supplemental Figure 1

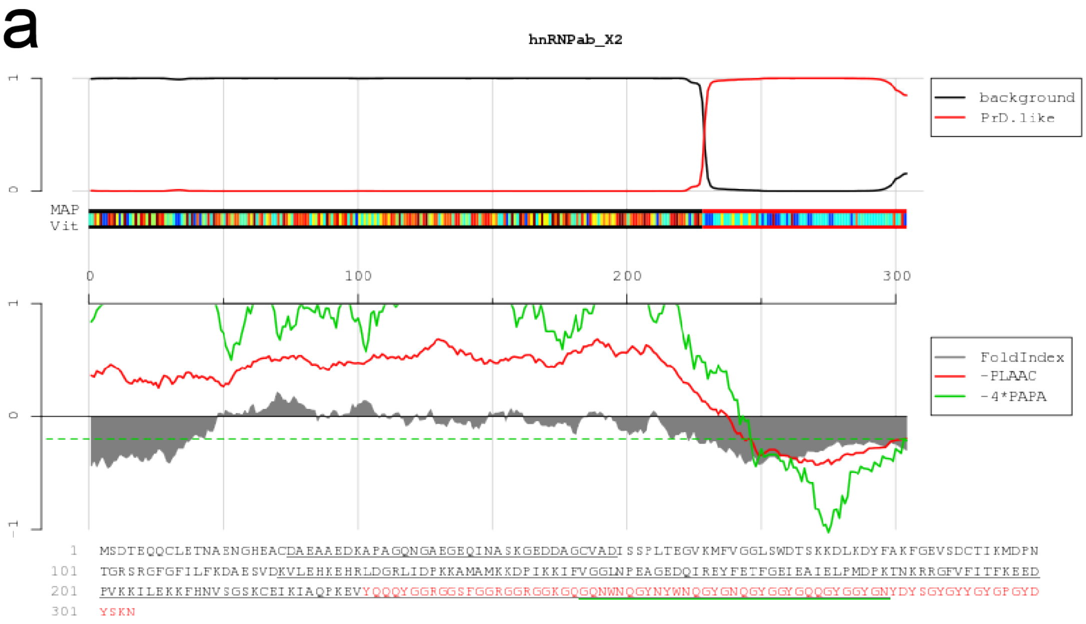


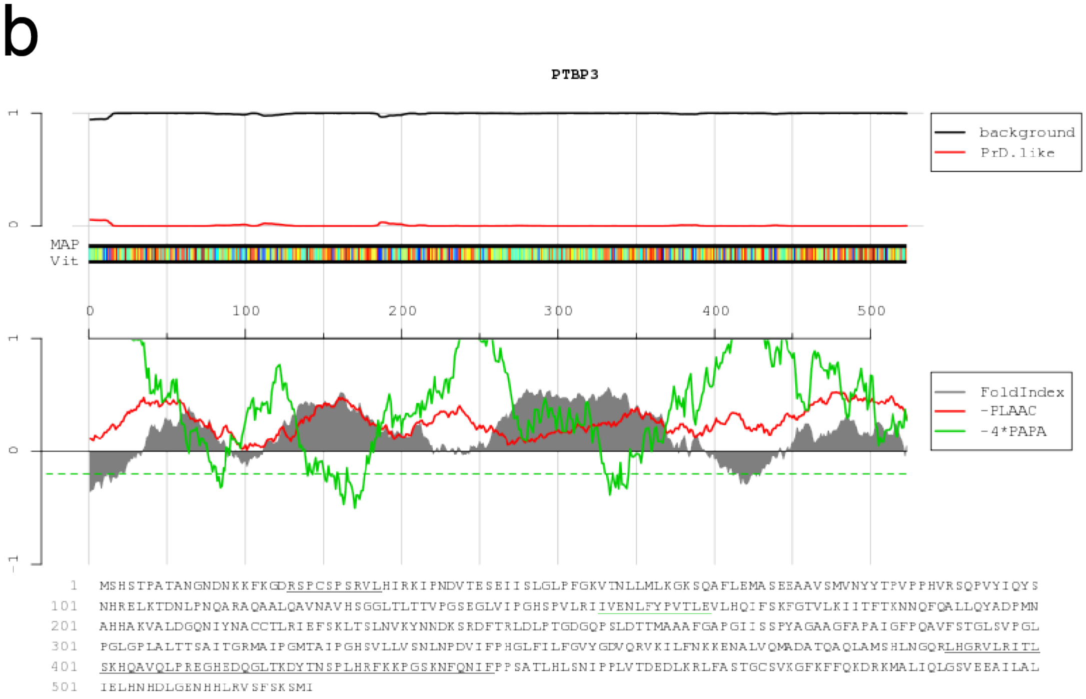


**Figure S1:** PLAAC Analysis of proteins. The protein sequences of **(a)** hnRNPAB X2 and **(b)** PTBP3 were analyzed using Prion-Like Amino Acid Composition (PLAAC)[1] to identify prion-like domains. The program identified a region at the C-terminus of hnRNPAB X2 as having a high degree of prion-like character (highlighted in red). PTBP3 was not found to have significant prion-like character.

**Supplemental Figure 2**

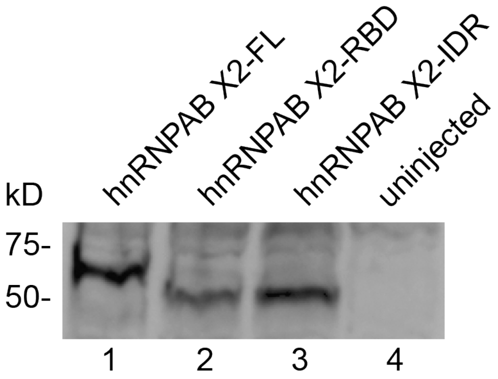


**Figure S2: Expression of hnRNPAB X2 domain constructs in oocytes.** Lysates were prepared from stage II oocytes injected with RNA transcribed from mCh-hnRNPAB X2 full-length (FL, lane 1), mCh-RBD (lane 2), mCh-IDR constructs, and uninjected control oocytes (lane 4). The constructs are diagrammed in Figure 2a. Immunoblot analysis was carried out using anti-mCh to detect the expressed domain constructs. The positions of molecular weight standards are indicated at the left.

**Supplemental Figure 3**

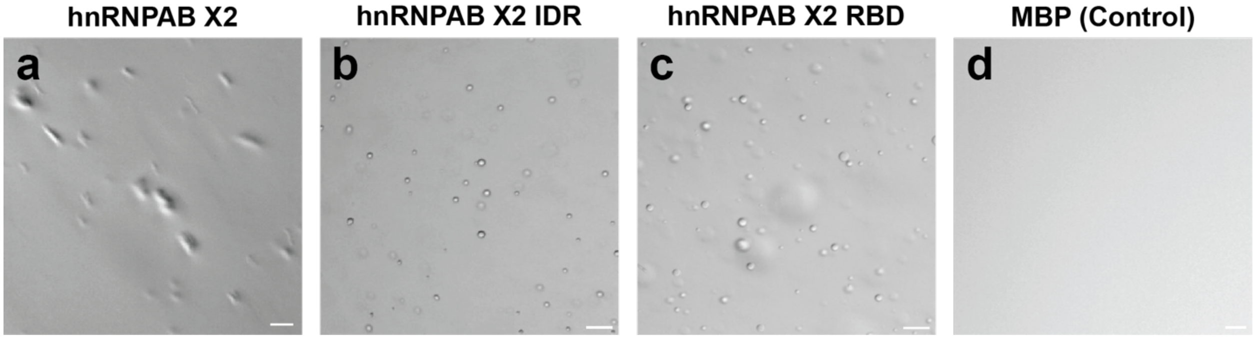


**Figure S3: In the presence of RNA, hnRNPAB X2 and its domains self-assemble and phase separate *in vitro* similarly as in absence of RNA (compare to Figure 3).** DIC micrographs of 50 µM MBP-fusions of (**a**) full-length hnRNPA2 X2, **(b)** IDR, (**c**) RBD, or the (**d**) control (free MBP) proteins in 20 mM NaPi, pH 7.4, 150 mM NaCl, 10% PEG and 0.25 mg/mL *Xenopus* beta globin RNA. Irregularly shaped assemblies were observed for hnRNPA2 X2 full-length while round droplets consistent with liquid-liquid phase separation (LLPS) were observed for X2 IDR and RBD. No phase separation was observed for free MBP (control). Scale bars: 20 µm.

**Supplemental Table 1**

| Protein Classification | % of L-body proteome | % of L-body IDR-containing proteins |
|---|---|---|
| No IDR | 53% | n/a |
| Non-Prion-Like IDR | 17% | 37% |
| Prion-Like IDR | 29% | 63% |

**Table S1:** Survey of L-body proteome for prion-like character. Using the L-body proteome[1] (*n*=86 proteins), L-body protein sequences available in Xenbase[2] were analyzed using PLAAC[1] to identify predicted IDRs and prion-like character. Proteins were rated as either having no IDR, a prion-like IDR, or a non-prion-like IDR, and percentages of the L-body proteome as a whole were generated based on these counts.

**Supplemental Table 2**

| Antibody | Source |
|---|---|
| Rabbit polyclonal anti-RFP | Abcam (ab62341) |
| Rabbit polyclonal anti-40LoVe | K. Czaplinski[3] |
| Goat anti-rabbit AF546 | Thermofisher (11010) |

**Table S2:** Antibodies used in this study. The name of the antibody and the antigen are specified on the left, and the source is shown on the right.

**Supplemental Table 3**

| Primer | Sequence |
|---|---|
| X2AB_FL_Fwd_GibsonCherry | GTACAAGAGATCTGATCATGCCATGGGGCCCATG TCCGACACCGAGCAGC |
| X2AB_RBD_Fwd_GibsonCherry | GTACAAGAGATCTGATCATGCCATGGGGCCCAAA ATGTTTGTTGGTGGCTTGAGCTG |

| | |
|---|---|
| X2AB_IDR_Fwd_GibsonCherry | GTACAAGAGATCTGATCATGCCATGGGGCCCAAAGAAGTGTATCAGCAACAGTATGGCG |
| X2AB_FL_Rev_GibsonCherrySpeI | GTTTAGTGGTAACCAGATCCTAGTCAGTCATCAGTTTTTACTGTAGTCATAGCCTGGTCC |
| X2AB_RBD_Rev_GibsonCherrySpeI | GTTTAGTGGTAACCAGATCCTAGTCAGTCATGGTTGTGCAATCTTTATCTCACACTTGC |
| hnRNPAB_PYNull_Cherry_Rev | GTTTAGTGGTAACCAGATCCTAGTCAGTCATCACTTGTAGTTATTCTGGTGGCTCCCAC |
| X2AB_FL_Fwd | TCCGACACCGAGCAGCAGTGTCTAGAA |
| X2AB_FL_Rev | GTTCTTACTGTAGTCATAGCCTGGTCCATATCCATAATAG |
| X2AB_RBD_Fwd | AAAATGTTTGTTGGTGGCTTGAGCTGG |
| X2AB_RBD_Rev | TGGTTGTGCAATCTTTATCTCACACTTG |
| X2AB_IDR_Fwd | AAAGAAGTGTATCAGCAACAGTATGGCGG |
| P3_IDR_Fwd | GTACAAGAGATCTGATCATGCCATGGGGCCCATGAGCCATTCCACTCCAGCTACAG |
| P3_IDR_Rev | CAGAGACAGAGACAGAGACAGAGAGATCATCGATTTGGAGAAAGAGACACG |
| XBM_Fwd | GAAAT<u>TAATACGACTCACTATAG</u>GGAGAGTTGAACTTGTAGCATCCAGCTCAGAATAAACGCTCAACTTTG |
| XBM_Rev | GGATCCACATGTAGGGTCTCT |
| Vg1_qPCR_Fwd | GGTATCTCCTCCTCCTGTCCCT |
| Vg1_qPCR_Rev | TGGGTGGATGTCATCGGAGT |
| CanAB_qPCR_Fwd | GGCTATTATGGATATGGACCAG |
| X2AB_qPCR_Fwd | ACAATTACTGGAACCAGGGCT |
| CanAB_qPCR_Rev | GGCTATTATGGATATGGACCAG |
| X2AB_qPCR_Rev | GTTTTTACTGTAGTCATAGCCTG |

**Table S3**: Primers used in this study. All Forward (Fwd) and Reverse (Rev) primers are listed from 5′ to 3′. The following abbreviations are used: X2AB (hnRNPAB X2), FL (full-length), P3 (PTBP3), XBM (*Xenopus* β globin), and CanAB (canonical hnRNPAB). XBM_Fwd contains a T7 promoter sequence (underlined).
